# Spatiotemporal Dissociation of Human Amygdala Response to Negative Affect

**DOI:** 10.64898/2026.08.18.745530

**Authors:** Ke Bo, Martin A. Lindquist, Peter J. Gianaros, Tor D. Wager

## Abstract

The amygdala is central to negative affect and a primary target of top-down regulation, yet its temporal dynamics remain poorly characterized. Most fMRI studies model stimulus-evoked amygdala activity with a canonical hemodynamic response function and collapse across subregions, potentially obscuring functional heterogeneity in time and space. We used finite impulse response modeling in two independent datasets (combined N = 358) to characterize blood oxygenation level-dependent responses during negative picture viewing and instructed cognitive reappraisal, complemented by data-driven clustering of voxelwise time courses. Across two probabilistic atlases, six of seven amygdala subregions showed prolonged activation to negative stimuli but differed in temporal profile. The centromedial amygdala showed sustained activation extending into the post-stimulus rating period, whereas laterobasal and superficial subregions peaked earlier during stimulus presentation. Reappraisal did not substantially alter response magnitude or shape, indicating that amygdala downregulation is not a reliable consequence of instructed regulation under these conditions. Data-driven analysis identified four clusters with distinct temporal profiles corresponding to laterobasal, lateral, superficial, and centromedial territories, converging with established anatomical parcellations while crossing some conventional boundaries. FIR modeling thus reveals a temporal architecture of amygdala responses to negative affective stimuli, characterized by early laterobasal and superficial responses followed by sustained centromedial activity.

****Significance Statement**:** Most human fMRI studies model the amygdala as a single region using a canonical hemodynamic response, potentially obscuring distinct temporal dynamics during affective processing. Across two independent cohorts, flexible time-course modeling revealed a reproducible sequence: laterobasal and superficial regions responded early to negative images, whereas centromedial activity persisted after image offset into the affective-evaluation period. Although cognitive reappraisal effectively reduced self-reported negative affect, it did not reliably alter these amygdala time courses. Data-driven clustering further recovered and extended this organization, showing that temporal response profiles align with, but are not fully constrained by, conventional anatomical boundaries.

## Introduction

The amygdala is a central hub in human emotion, consistently activated across a wide range of affective paradigms, including processing emotional scenes, faces, films, and odors ^1–3^. In studies of emotion regulation, it has been treated as the primary target of top-down control, with reduced amygdala activity widely interpreted as evidence of successful regulation ^4–6^. Yet despite its robust engagement by affective stimuli, evidence that emotion regulation reliably modulates amygdala activity has been inconsistent. The regulatory effect of cognitive reappraisal on amygdala activity has been reported in some studies ^4,5^ but found to be absent in others ^7,8^. Such inconsistencies challenge a unified understanding of the amygdala’s role in affective processing and point to unresolved questions about both its functional organization and the conditions under which it is susceptible to regulatory influence.

Two fundamental limitations may contribute to these mixed findings. The first concerns how amygdala activity is measured. Most fMRI analyses rely on the canonical hemodynamic response function (HRF), derived primarily from neocortical regions such as visual and motor cortex ^9^. However, evidence from high-resolution 7T fMRI suggests that subcortical visual structures, such as the LGN and superior colliculus, exhibit shorter time-to-peak and narrower hemodynamic response profiles than primary visual cortex, indicating that canonical HRF assumptions may not fully capture subcortical BOLD dynamics ^10^. When the assumed response shape mismatches the true neural signal, standard modeling approaches will mischaracterize both the timing and magnitude of amygdala responses, potentially contributing to inconsistencies across studies^11,12^.

The second limitation is the frequent treatment of the amygdala as a unitary region of interest. In reality, the amygdala is a heterogeneous structure composed of multiple nuclei with distinct cytoarchitecture, connectivity, and functional roles ^13,14^. The laterobasal (LB) nucleus of the amygdala receives extensive cortical and thalamic input and is thought to serve as a sensory gate for rapid evaluation of emotional significance ^15–17^. The centromedial (CM) nucleus serves as the principal output nucleus of the amygdala, projecting to brainstem, hypothalamic, and striatal targets that coordinate autonomic and behavioral responses to salient and threatening stimuli ^16,18,19^. The superficial amygdala is implicated in processing socially relevant and affective cues, particularly in the context of olfactory and face processing ^20,21^. These findings highlight the importance of studying amygdala subnuclei separately, rather than collapsing across them. In sum, regional heterogeneity within the amygdala and temporal variation in subcortical hemodynamic responses together suggest that amygdala function is likely organized in both space and time, yet few studies have tested whether anatomically distinct subregions exhibit dissociable response time courses that could provide a more accurate account of affective processing.

A powerful approach to characterize such temporal dynamics is the Finite Impulse Response (FIR) model, which estimates the BOLD signal as a freely varying time course at each time point following stimulus onset, without assuming a fixed canonical shape ^22^. FIR analyses have been used to reveal onset latency, peak timing, and response duration differences across brain regions that canonical models cannot capture ^11,23^. In the amygdala specifically, where responses may deviate substantially from cortical hemodynamics and where trial structure often confounds temporally adjacent processes, FIR modeling is especially well suited to recovering the full temporal profile of activity. However, this flexibility comes at a cost: because FIR modeling estimates a separate coefficient at each post-stimulus time point, it is less statistically efficient than more constrained HRF models and therefore particularly benefits from large, well-powered datasets when the goal is to compare time courses across small subcortical subregions and conditions.

In the current study, we applied FIR modeling to fMRI data from 358 healthy adults across two independent datasets, providing a well-powered framework for stable estimation of BOLD time courses. Participants completed a picture-viewing paradigm that included both passive viewing of negative and neutral images and a cognitive reappraisal condition. We defined amygdala subregions using two independent probabilistic atlases, the Amunts et al. atlas ^24^ and the Nacewicz et al. atlas ^25^, to ensure that our findings were not dependent on any single parcellation scheme. Related probabilistic masks derived from these atlases have been widely used in prior human neuroimaging studies of amygdala subregions and extended-amygdala circuitry ^15,26,27^. We asked two central questions: first, do amygdala subregions exhibit distinct temporal dynamics in response to negative emotional stimuli; and second, does cognitive reappraisal differentially modulate these temporal profiles across subregions? We further applied a data-driven clustering approach to group amygdala voxels by their FIR-derived temporal profiles, allowing us to assess whether functionally defined temporal clusters correspond to known anatomical boundaries. Together, these analyses provide a fine-grained characterization of the temporal architecture of amygdala function during human negative affect.

## Results

### Behavioral manipulation checks confirmed successful affect induction and regulation

Behavioral manipulation checks from these cohorts have been reported previously ^7^. Negative-affect ratings were substantially higher during Look Negative than Look Neutral trials, confirming successful affect induction (t = 63.8, P < 0.001, Cohen’s d = 3.37). Ratings were also lower during Regulate Negative than Look Negative trials, confirming successful cognitive reappraisal (t = −17.5, P < 0.001, d = 0.92). Eighty-one percent of participants showed a regulation effect in the same direction as the group-level effect. Thus, participants successfully reduced their reported negative affect despite the absence of reliable reappraisal-related modulation of amygdala BOLD time courses.

### Amygdala subregions engage early and sustain activation to negative stimuli

As shown in Figure 1, each trial comprised a 2-s cue period, a 7-s stimulus period, and a 4-s rating period. With a TR of 2 s, these correspond to approximately 1, 3.5, and 2 TRs, respectively. Accounting for the hemodynamic delay of the BOLD response (∼3 TRs to peak), the cue, stimulus, and rating periods are expected to be most strongly represented at approximately TRs 3–4, 4–7, and 8–10. To characterize the temporal dynamics of amygdala subregions during affective processing, we estimated FIR time courses for each subregion in response to the IAPS images across 15 TRs, a window chosen to capture the full trial epoch including the delayed hemodynamic response.

**Figure 1.**
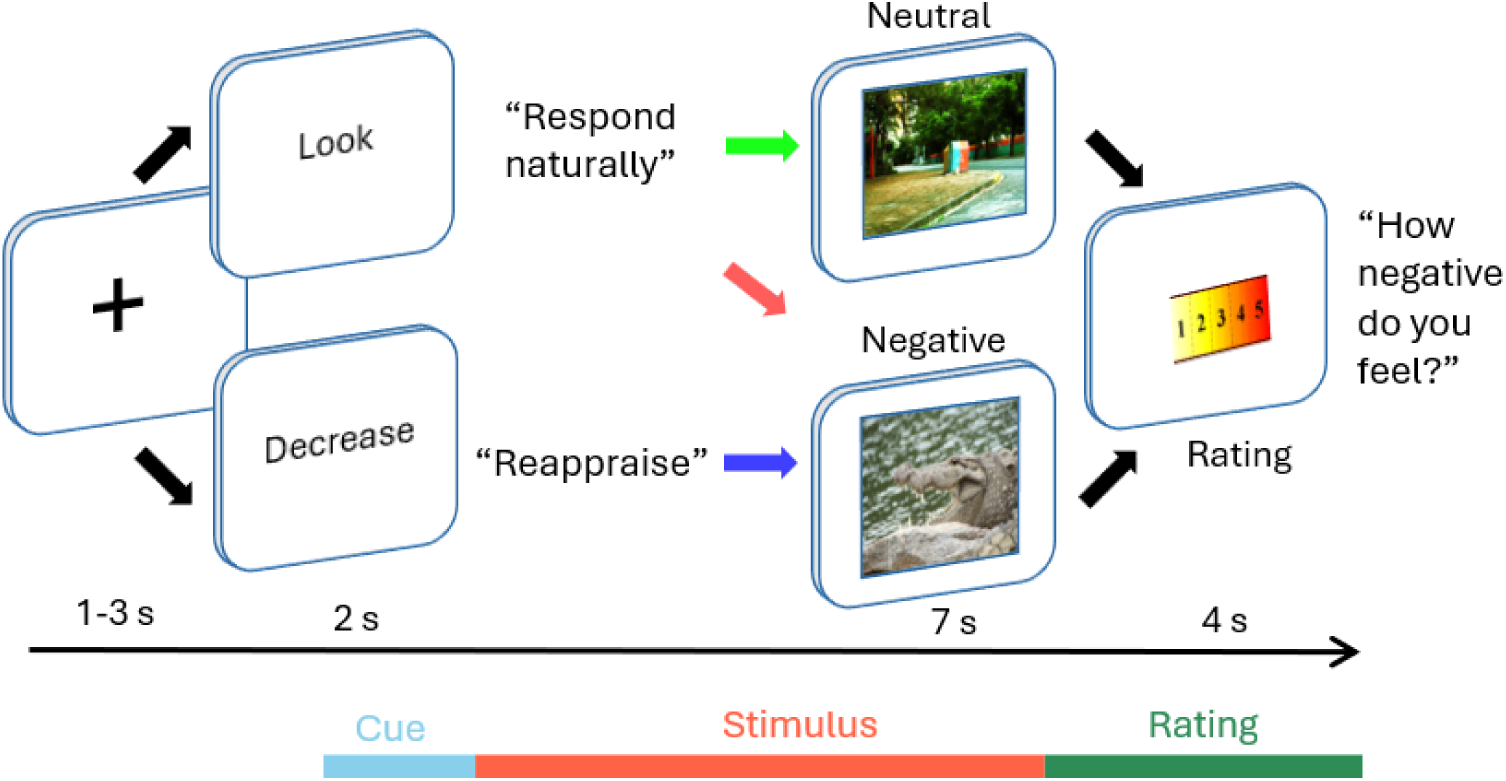
The overall trial structure: Trials followed a fixed sequence consisting of a 2-s appraisal cue (“Look” or “Decrease”), a 7-s image-viewing period, and a 4-s negative-affect rating. Depending on the condition, participants either viewed neutral images passively (“Look neutral”), viewed aversive images passively (“Look negative”), or used cognitive reappraisal to downregulate negative affect in response to aversive images (“Regulate negative”). With TR of 2s, accounting for the hemodynamic delay of the BOLD response (∼3 TRs to peak), the cue, stimulus, and rating periods are expected to be most strongly represented at approximately TRs 3–4, 4–7, and 8–10.

The FIR-estimated BOLD time courses for the two atlases are shown in Figure 2. First, we tested whether emotion pictures evoke reliable activation compared to baseline. In the Amunts atlas, all three subregions exceeded baseline significantly at TR 4. The superficial amygdala (SF) and laterobasal amygdala (LB) sustained above-baseline activation through TR 8, peaking at TR 5 within the stimulus window (SF: t(357) = 7.18; LB: t(357) = 6.65, p < 0.05, FDR corrected). The centromedial amygdala (CM), by contrast, showed a temporally distinct pattern: although it also became significant at TR 4, its response continued to rise after picture offset, peaking at TR 8 (t(357) = 13.15, p < 0.05, FDR corrected), which likely falls within the rating period BOLD window rather than the stimulus period. This suggests that CM activation reflects not only perceptual processing of the negative image but continued engagement during the subsequent rating period. In the Nacewicz atlas, CoAHA and BLBM similarly rose above baseline at TR 4 and peaked at TR 5, within the stimulus window (CoAHA: t(357) = 9.88; BLBM: t(357) = 6.86, p < 0.05, FDR corrected). CeMe mirrored CM with activation spanning TRs 4–9 and a peak at TR 7 (t(357) = 8.06, p < 0.05, FDR corrected), again consistent with sustained engagement into the rating period. The lateral nucleus (LA) showed the most limited response, reaching significance only at TR 7 (t(357) = 2.63, p < 0.05, FDR corrected).

**Figure 2.**
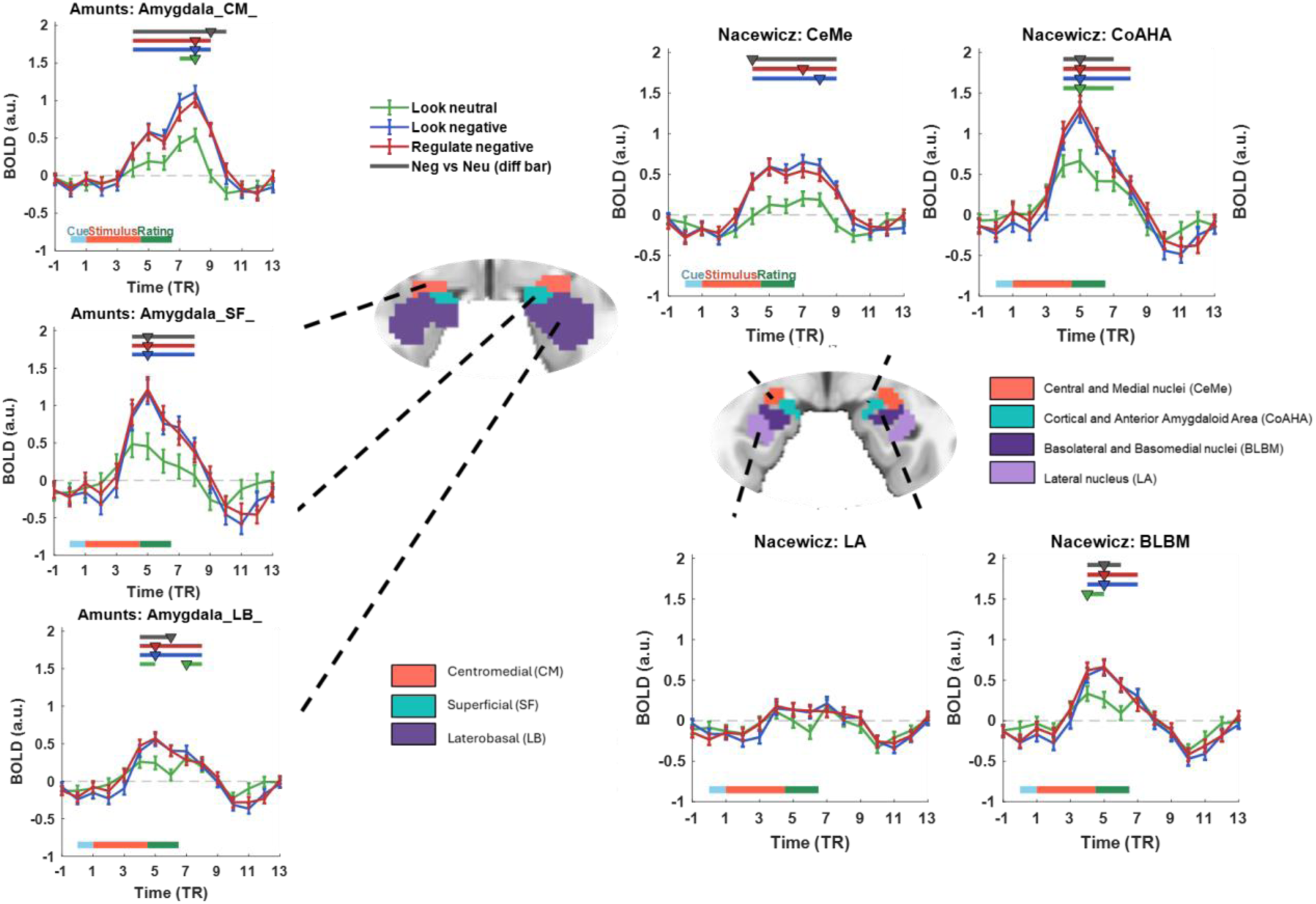
Temporal dynamics of amygdala subregions during emotional picture viewing and reappraisal, estimated by finite impulse response (FIR) modeling. BOLD responses were modeled with an FIR basis set spanning 30 s (15 TRs; TR = 2 s) for each of three task conditions. Curves show the group-mean FIR response (± SEM) for Look Neutral (green), Look Negative (blue), and Regulate Negative (red). The time axis is referenced to cue onset: TR 0 = cue onset and TR 1 = stimulus (picture) onset. The colored bar beneath each time course marks the trial structure: cue (0–1 TR), stimulus presentation (1–4.5 TR), and rating (4.5–6.5 TR). Horizontal bars above each time course denote intervals of statistically significant activation, identified by per-time point one-sample t-tests against baseline (paired t-tests for the condition contrast), FDR-corrected across the 15 TRs (Benjamini-Hochberg, q < .05); only positive effects spanning ≥2 consecutive significant TRs are shown. Bar colors match the curves: green = Look Neutral vs. baseline, blue = Look Negative vs. baseline, red = Regulate Negative vs. baseline, and grey = Look Negative vs. Look Neutral. The inverted triangle (▾) on each bar marks the peak TR of that effect. Error bars indicate the standard error of the mean (SEM) across participants. These plotting conventions are used consistently across all time-series figures. **Left:** Amunts atlas: centromedial (CM), superficial (SF), and laterobasal (LB) amygdala. **Right:** Nacewicz atlas: central and medial nuclei (CeMe), cortical and anterior amygdaloid area (CoAHA), basolateral and basomedial nuclei (BLBM), and the lateral nucleus (LA). The central insets show the anatomical location of each subregion on a coronal template.

Subsequently, to isolate emotion-specific activation, we compared look-negative against look-neutral evoked-response time course using paired t-tests at each TR (FDR corrected). Across both atlases, negative images produced reliably greater activation than neutral images beginning at TR 4 and sustained across multiple consecutive TRs (Figure 2). In the Amunts atlas, CM showed the longest sustained differentiation, spanning TRs 4–10 (7 TRs, 14 seconds; peak t(357) = 7.93 at TR 9, p < 0.05, FDR corrected). Critically, this sustained window extended into the rating period, suggesting that CM differentiates negative from neutral content even after picture offset. SF differentiated from TR 4 to TR 8 (peak TR 5: t(357) = 6.48). LB showed a window of differentiation at TRs 4–6 (peak TR 6: t(357) = 5.14). In the Nacewicz atlas, CeMe again showed the most sustained differentiation, spanning TRs 4–9 (peak TR 4: t(357) = 6.77). CoAHA differentiated across TRs 4–7 and BLBM across TRs 4–6 (CoAHA peak TR 5: t(357) = 6.27; BLBM peak TR 5: t(357) = 5.51). LA showed no sustained differentiation, defined as at least two consecutive significant TRs. Given the trial structure and hemodynamic delay, the BOLD responses corresponding most closely to the cue, stimulus, and rating periods were expected at approximately TRs 3–4, 4–7.5, and 7.5–9.5, respectively.

### The centromedial amygdala uniquely sustains activation at late TRs

As shown in Figure 3, between-region comparisons revealed a consistent pattern in which CM (Amunts) and CeMe (Nacewicz) diverged from other subregions at late TRs. In the Amunts atlas, CM was significantly less activated than SF at TRs 4–5 but substantially exceeded SF from TR 7 onward (TRs 7–11; peak TR 9: t(357) = 6.43, p < 0.05, FDR corrected). CM also exceeded LB across TRs 7–10, with a particularly large difference at TR 8 (t(357) = 11.54, p < .05, FDR-corrected). In the Nacewicz atlas, CeMe was similarly smaller than CoAHA at the early peak (TRs 4–6), but exceeded it from TR 8 onward (TRs 8–11; peak TR 9: t(357) = 4.90). CeMe also exceeded LA (TRs 4–9; peak TR 8: t(357) = 6.55) and BLBM (TRs 7–11; peak TR 8: t(357) = 8.33) across a broad late window. Together, these results indicate that CM and CeMe are not simply more activated overall, but exhibit a temporally distinct, late-sustained elevation that other subregions do not show.

**Figure 3.**
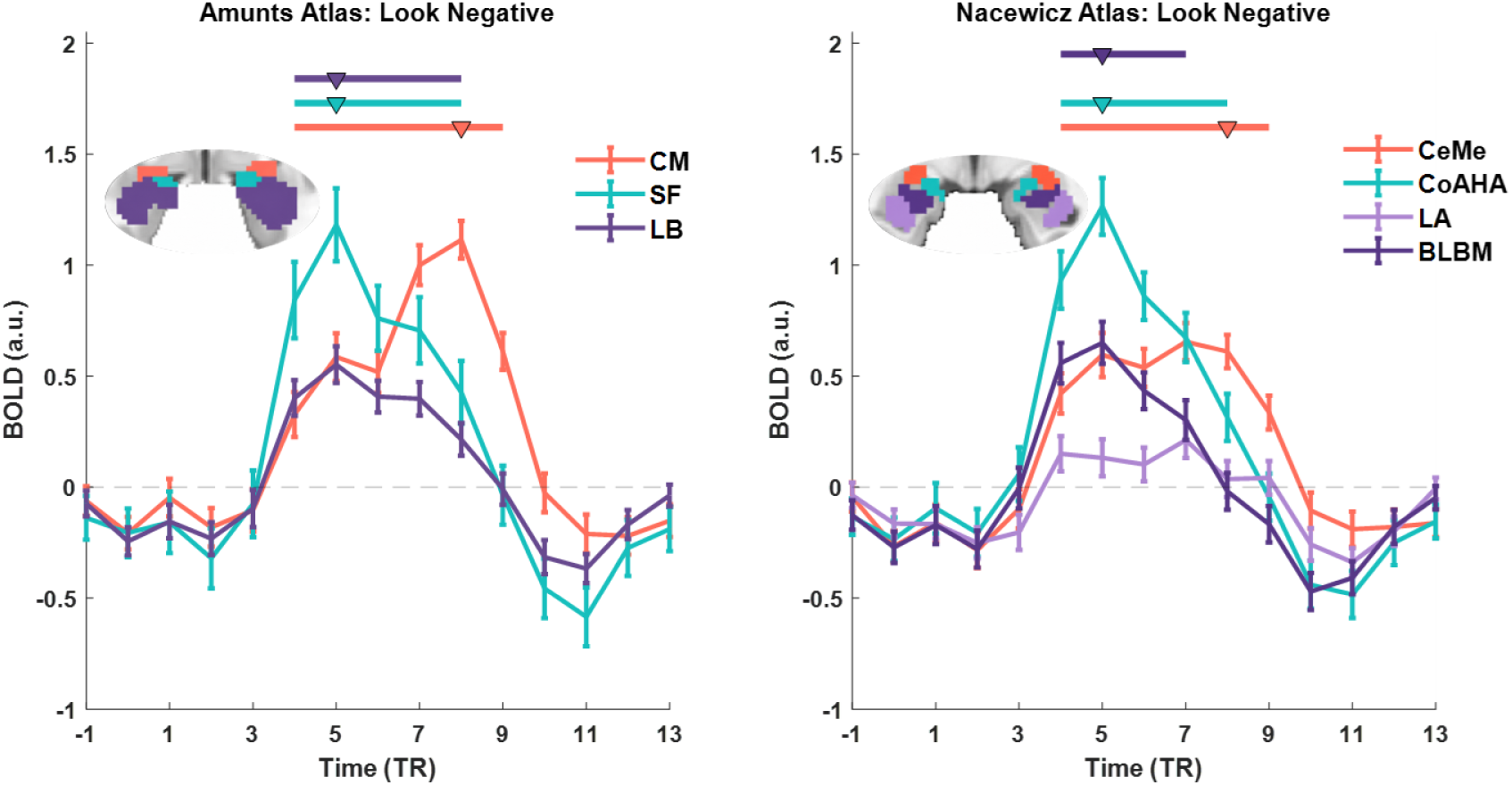
Temporal dynamics of amygdala subnuclei based on FIR analysis, compared directly on the same axis using the Look Negative condition. A) Amunts atlas, including the Centromedial (CM), Laterobasal (LB), and Superficial (SF) nuclei. B) Nacewicz atlas, including the Central and Medial nuclei (CeMe), Cortical and Anterior Amygdaloid Area (CoAHA), Basolateral and Basomedial nuclei (BLBM), and Lateral Nucleus (LA).

### Instructed cognitive reappraisal produces no detectable modulation of amygdala evoked-response time courses

We next asked whether cognitive reappraisal modulated amygdala temporal dynamics of activation. We compared look-negative and regulate-negative evoked-response time courses at each TR (paired t-tests, FDR corrected). After FDR correction, down-regulation effects were largely absent (Look negative > Regulate negative) across both atlases. At an uncorrected level, CM showed a numerical trend toward look-negative > regulate at TR 7 (t(357) = 2.68, *d* = 0.14, p = 0.008), though this did not survive FDR correction. A two-sided JZS Bayes factor analysis using the default zero-centered Cauchy prior on the standardized effect size (scale r 0.707) was conducted at this time point. The analysis yielded BF₁₀ = 2.02 at this time point, indicating only weak evidence for a difference relative to the null hypothesis. Notably, this effect occurred near the end of the stimulus period and the beginning of the rating period considering hemodynamic delay. Taken together, these results provide no reliable evidence that cognitive reappraisal downregulates amygdala BOLD responses at the time-course level. Any potential effect was small, restricted to the centromedial amygdala, and not robust to correction for multiple comparisons.

### Lateralization analysis

Prior neuroimaging studies have reported left-dominant amygdala activation during emotional processing ^29,30^, though whether this asymmetry extends to specific subregions and trial periods remains unclear. To examine hemispheric asymmetry, we re-extracted evoked-response time courses separately for left and right hemisphere subregions and applied paired t-tests at each TR (FDR-corrected, p < .05), along with a laterality index (LI = [Left − Right] / [|Left| + |Right|]) at the group peak TR for the look negative condition.

As shown in figure 4, lateralization effects were region-specific and most pronounced during look negative and regulate conditions. In the Amunts atlas, the centromedial nucleus (CM) showed consistent left-hemisphere dominance: left CM exceeded right CM at TRs 7–10 during look negative (t(357) = 3.91–5.15, p < .05) and at TRs 7–9 during regulate (t(357) = 5.38–6.43, p < .05), but not during look neutral. The LI confirmed significant leftward asymmetry at the peak response (LI = 0.187 ± 0.032, t(357) = 5.85, p < .001). The superficial nucleus (SF) showed a smaller but significant leftward LI (LI = 0.078 ± 0.036, t(357) = 2.18, p = .030), while the laterobasal nucleus (LB) showed no lateralization (LI = 0.001, p = .983). In the Nacewicz atlas, the CoAHA region showed left-greater-than-right differences at late TRs during look negative (TRs 8–9; t(357) = 3.14–3.71, p < .05, FDR corrected) and regulate (TR 8; t(357) = 4.14, p < .05, FDR corrected), though its peak-TR LI was not significant (p = .902). CeMe showed significant leftward lateralization at only one late TR after FDR correction during look negative (TR 8; t(357) = 3.43, p < .05, FDR corrected). No other Nacewicz regions were significantly lateralized.

**Figure 4.**
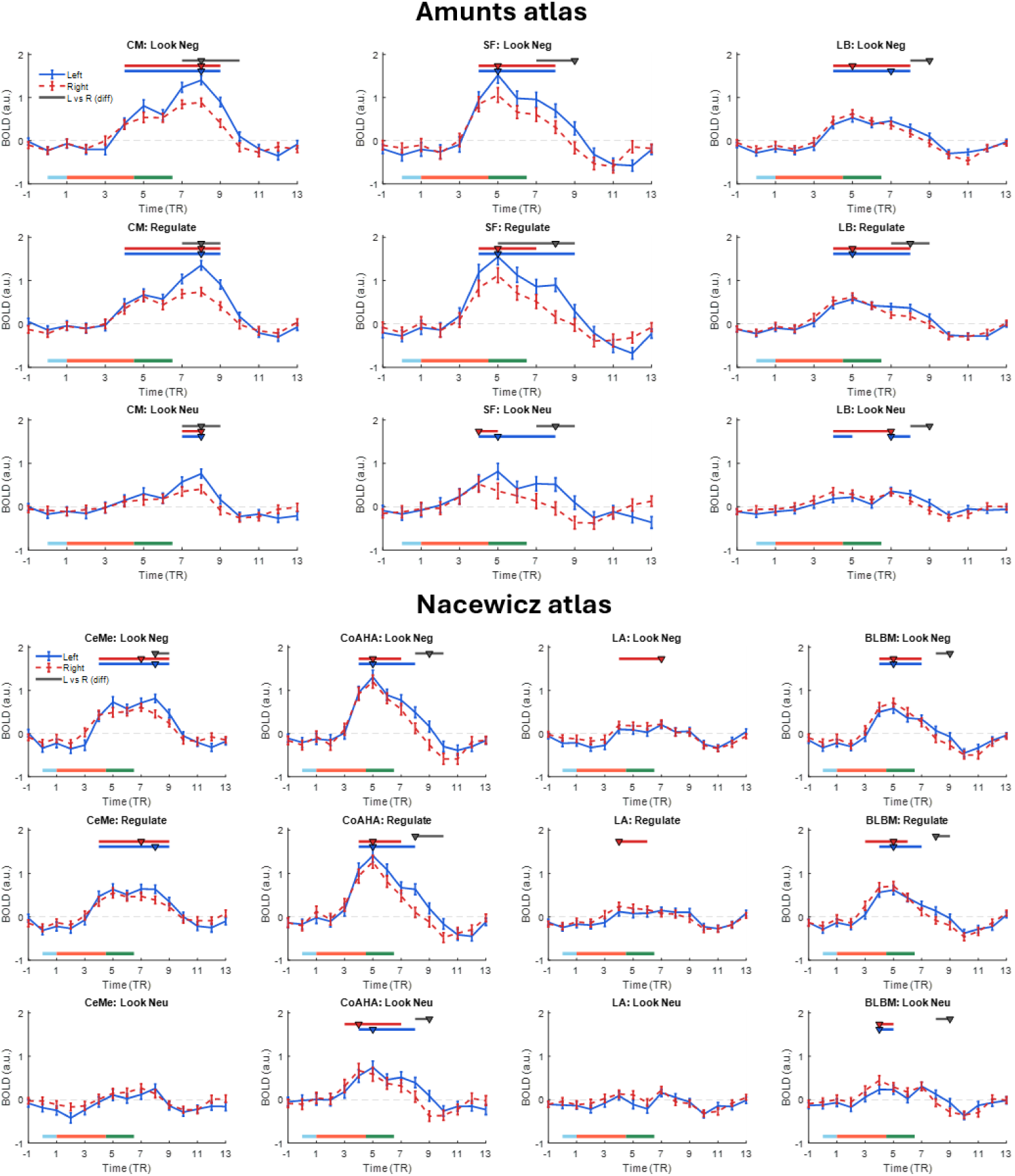
Hemispheric lateralization of FIR-estimated BOLD responses. Each panel shows group-mean FIR time courses (±SEM) for left (solid blue) and right (dashed red) hemisphere subregions across three conditions: look negative (top), regulate (middle), and look neutral (bottom). The x-axis represents time post-cue onset in TRs (TR = 2s); the y-axis represents BOLD amplitude in arbitrary units. **Top)** Amunts atlas: Columns correspond to centromedial (CM), superficial (SF), and laterobasal (LB) subregions. **Bottom)** Nacewicz atlas: Columns correspond to central and medial nuclei (CeMe), cortical and anterior amygdala area (CoAHA), lateral nucleus (LA), and basolateral and basomedial nuclei (BLBM).

Overall, hemispheric asymmetry was selective rather than global. It was observed most consistently in the centromedial amygdala during late TRs, likely corresponding to the rating period, during negative affective processing and reappraisal. This is the same region that showed the most distinctive delayed temporal profile in the bilateral analysis.

### Data-driven clustering based on temporal profiles of amygdala voxels

The subregional analyses above relied on anatomical definitions imposed a priori. To test whether a similar organization emerged directly from evoked affective response time courses, we clustered all 958 amygdala voxels according to their 45-dimensional FIR feature vectors (15 TRs × 3 conditions) using k-means clustering, without incorporating anatomical labels. A permutation-based model-selection analysis identified *k* = 4 as the optimal solution (see Methods). The four clusters contained 255, 474, 140, and 89 voxels. Notably, each cluster exhibited a distinct temporal response profile (Figure 5B) and showed significant spatial correspondence with subregions in both the Nacewicz and Amunts anatomical parcellations.

**Figure 5.**
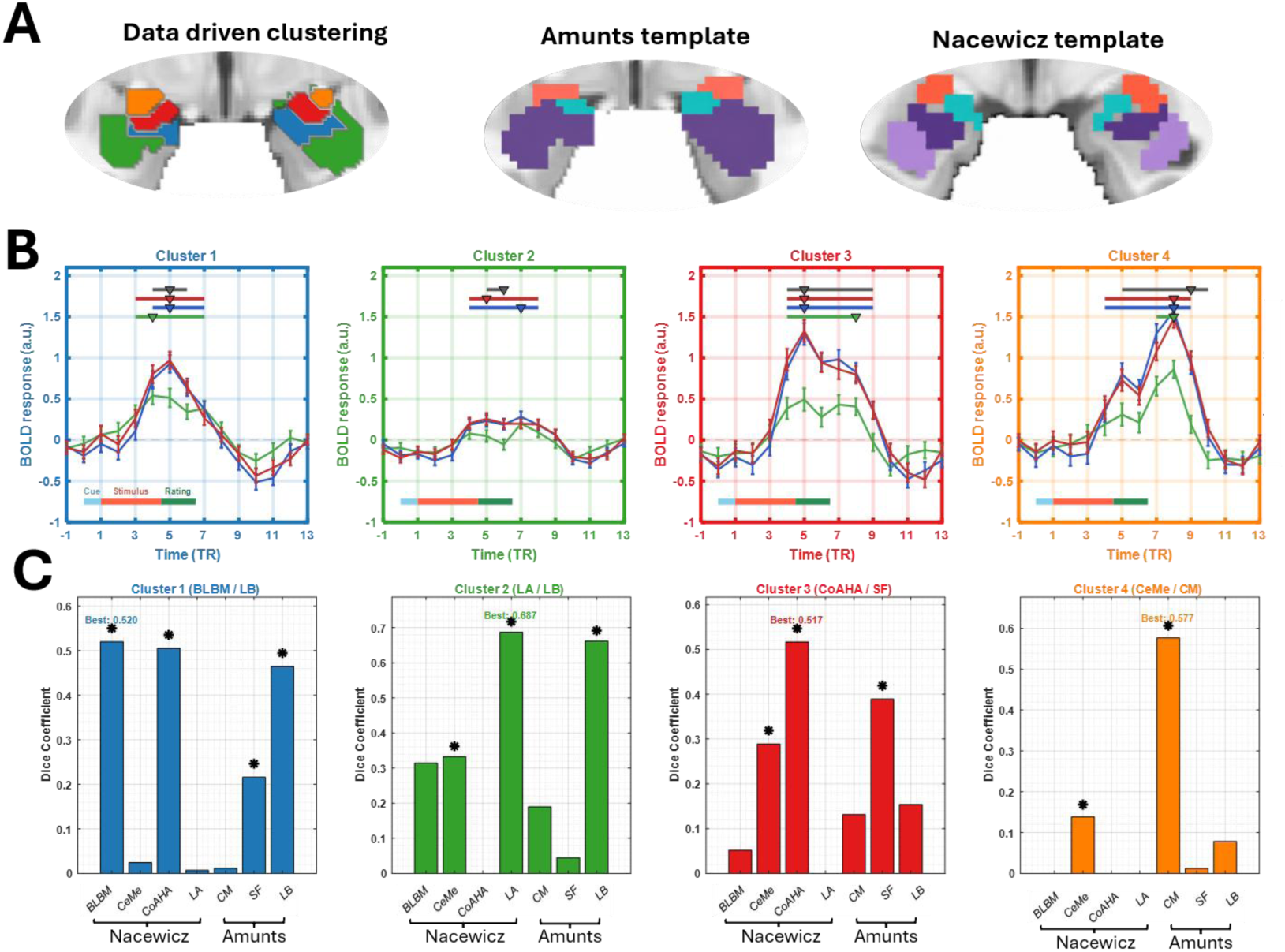
Data-driven clustering across the combined amygdala masks from the Amunts and Nacewicz atlases. **A)** Spatial locations of k-means-derived clusters of amygdala voxels, shown in comparison with the Amunts and Nacewicz anatomical atlases. **B)** Mean FIR time series for each data-driven cluster. **C)** Dice coefficients showing the spatial overlap between data-driven clusters and anatomical amygdala subregions. Asterisks indicate overlaps that were significant relative to a permutation null distribution, with multiple comparisons corrected across the full cluster-by-region matrix using Benjamini-Hochberg FDR at *q* < 0.05.

To compare the spatial distributions of the data-driven clusters with the anatomical parcellations, we first quantified global agreement using the adjusted Rand index. Agreement was 0.276 for the Nacewicz parcellation and 0.161 for the Amunts parcellation. Both values were significantly greater than expected by chance based on 10,000 permutations (both permutation *p* < .0001). Thus, although the data-driven and anatomical boundaries were not identical, the organization of amygdala voxels based on their temporal response profiles showed significant, nonrandom correspondence with anatomical organization.

We next characterized the temporal profile and anatomical distribution of each cluster (Figure 5). Cluster 1 exhibited a relatively rapid and transient response, with responses to negative pictures peaking at TR 5 and greater activation for Look Negative than Look Neutral during the early response period. The response subsequently returned toward baseline without a prominent late second peak. In the Nacewicz parcellation, Cluster 1 overlapped significantly with both BLBM (Dice = 0.520) and CoAHA (Dice = 0.505), with nearly equivalent overlap across the two regions. In the Amunts parcellation, it corresponded most strongly to LB (Dice = 0.465). Thus, this early and transient response profile was not restricted to a single nucleus but crossed the boundary between the deep basal or laterobasal territory and the neighboring cortical–amygdalohippocampal territory. Its correspondence with Amunts LB nevertheless placed much of the cluster within the broader laterobasal division.

Cluster 2 exhibited the weakest evoked response of the four clusters, with relatively modest differentiation between negative and neutral pictures. Anatomically, it matched LA in the Nacewicz parcellation (Dice = 0.687) and LB in the Amunts parcellation (Dice = 0.662). The convergence across these parcellations localized this comparatively low-amplitude affective response profile to the lateral portion of the broader laterobasal complex.

Cluster 3 showed a robust early response that peaked at TR 5 and remained elevated during the subsequent portion of the trial. Look Negative exceeded Look Neutral across an extended early-to-middle response period. Cluster 3 corresponded most strongly to CoAHA in the Nacewicz parcellation (Dice = 0.517), but it also showed significant secondary overlap with CeMe (Dice = 0.289). In the Amunts parcellation, its strongest correspondence was with SF (Dice = 0.389). These convergent results indicate that the sustained affective response was centered in the superficial ventromedial and amygdalohippocampal territory but extended into the adjacent centromedial amygdala.

In contrast, Cluster 4 displayed the most delayed temporal profile. Its responses continued to increase later in the trial and reached their maximum at TR 8, with Look Negative exceeding Look Neutral across the late response period. Cluster 4 corresponded significantly with CeMe in the Nacewicz parcellation (Dice = 0.139) and showed a much stronger correspondence with CM in the Amunts parcellation (Dice = 0.577). Despite the difference in overlap magnitude across atlases, both parcellations localized this delayed response profile to the dorsal-centromedial amygdala. This data-driven result independently recovered the delayed centromedial response observed in the atlas-based analyses.

None of the four clusters showed a significant difference between Regulate Negative and Look Negative after FDR correction, indicating that the distinctions among clusters primarily reflected differences in the timing and magnitude of evoked affective responses rather than reliable effects of instructed down-regulation.

It should be noted that the magnitude of the Dice coefficient does not map directly onto statistical significance: because the permutation null for each pair depends on the sizes of both the cluster and the region, some smaller Dice values reach significance while some larger ones do not (See Methods for details).

### Replication and differences across two independent cohorts

To verify that the results were replicated across two independent datasets, we repeated all analyses separately within each dataset (AHAB cohort, n = 182; PIP cohort, n = 176; Supplementary Material). The central findings reproduced in both cohorts. The delayed centromedial response again distinguished CM/CeMe from the earlier-peaking subregions (peak at TRs 7-8 vs. TR 5), with Look Negative evoked-response time courses correlating across cohorts at r > 0.80 for all atlas subregions except for LA (r = 0.57, see detailed results in Supplementary Figure 1). The leftward lateralization of the centromedial subregion was also robust (Look Negative laterality index: AHAB = +0.194, PIP = +0.162, both p < .001, see detailed results in Supplementary Figure 2).

The data-driven clustering also replicated: the four-cluster solution corresponded to both atlases well above chance in each sample (ARI = 0.258 and 0.210 for Nacewicz and Amunts in AHAB; 0.214 and 0.132 in PIP; all permutation p ≤ .0002), and the partitions themselves reproduced across cohorts (ARI = 0.369, p < .001; matched-cluster Dice = 0.62–0.79). The cluster overlapping the centromedial subregion did so at an identical Dice of 0.578 in both samples, again exhibited the most sustained late differentiation, and was matched across cohorts at the highest overlap of any cluster pair (Dice = 0.791; Look Negative time course r = 0.971). Effects outside the centromedial subregion were more variable: the superficial-subregion lateralization and the negative-versus-neutral differentiation of the basolateral and cortical subregions reached significance in AHAB but were attenuated in PIP, and no Nacewicz subregion was reliably lateralized in both cohorts (Supplementary Figure 3). No down-regulation effect replicated across cohorts: isolated significant TRs in one cohort did not recur in the other, and no region showed a consistent reappraisal effect across samples. Thus the temporally distinct, left-lateralized centromedial response is robust across independent samples, whereas weaker subregional effects should be interpreted with corresponding caution.

## Discussion

Using finite impulse response modeling in two large independent fMRI datasets, we characterized the temporal dynamics of the amygdala during negative affect and cognitive reappraisal. Three findings emerged. First, the amygdala sustains activation to negative stimuli well beyond stimulus offset, with no meaningful anticipatory response during the cue period.

Second, and most importantly, this sustained activation has internal structure: the centromedial nucleus shows a distinct late response that persists into the post-stimulus evaluation period, whereas laterobasal and superficial subregions show an earlier, more transient profile. Third, data-driven clustering of voxelwise temporal dynamics, without anatomical priors, recovers spatial divisions that converge with this organization. Under the instructed-reappraisal paradigm used here, none of these temporal profiles within amygdala was substantially modulated by cognitive regulation.

Some general features of the amygdala evoked-response time courses are worth discussing. Across all subregions and conditions, no significant deflection appeared at TR 3, which most likely indexes cue-related processing after accounting for the 0–2 s cue period and hemodynamic delay. This suggests that within picture-based affective paradigms, the amygdala is engaged primarily by the affective stimulus itself rather than by predictive cues or preparatory regulatory signals, consistent with prior work emphasizing its role in stimulus-locked rather than anticipatory processing ^31,32^.

Beyond the general temporal profile, we observed a clear functional dissociation across amygdala subregions. The early LB/SF response is consistent with the known anatomical position of these nuclei as the principal input stage of the amygdala. The laterobasal complex receives highly processed sensory information from temporal and parietal cortex and is considered the locus at which perceptual features are integrated with emotional significance ^17,27,33^. An early peak therefore aligns with a role in rapid appraisal of incoming stimuli and integrating sensory input. The superficial group, which has been implicated in the evaluation of socially and biologically relevant cues^20,33^, showed a similar early profile, suggesting a similar evaluation role of social information in the emotion pictures, since many of the images contained human faces and bodies.

The CM showed a distinct dual-peak temporal profile: an early peak coincident with stimulus presentation, followed by a delayed second peak at 8–9 TR during the rating period when the picture was no longer on screen. Anatomically, CM serves as the principal output stage of the amygdala, receiving input from BL and projecting to brainstem and hypothalamic targets that mediate autonomic and behavioral responses ^13,17^. Primate single-unit recordings further suggest that, while BL neurons evaluate the emotional significance of stimuli, CM neurons allocate attention to significant stimuli and initiate situation-appropriate autonomic responses ^34^. The early CM peak in our data thus aligns with this attentional allocation role. Regarding the late CM response, we interpret it as reflecting sustained affective evaluation, a process by which the affective value of the just-presented stimulus is maintained, integrated, and elaborated to inform an internal judgment in the absence of ongoing sensory input. This interpretation is consistent with prior evidence that amygdala activity can persist for several seconds after stimulus offset to support ongoing elaboration of affective information ^35^, and with appraisal-based accounts in which the amygdala flexibly tunes its activity to the motivational and evaluative significance of stimuli for the perceiver’s current goals ^36–38^. In summary, this late CM response may reflect a new role of the centromedial nucleus in emotion appraisal beyond sensory integration.

The data-driven clustering analysis further supported this organization. Grouping voxels solely by their temporal profiles produced spatially coherent clusters that moderately corresponded to cytoarchitectonic subdivisions, indicating that amygdala temporal dynamics are constrained by, but not reducible to, anatomical architecture. The clusters captured a spectrum of response profiles: an early and transient response spanning laterobasal and amygdalohippocampal territories, a comparatively weak response localized to the lateral laterobasal complex, an early but sustained response centered in superficial and amygdalohippocampal regions, and a delayed response localized to the dorsal centromedial amygdala. Thus, functional temporal boundaries did not map perfectly onto individual anatomical nuclei, but the data-driven solution independently recovered the principal atlas-based distinction between earlier responses in laterobasal and superficial territories and delayed activity in the centromedial amygdala. Temporal-profile-based parcellation may therefore provide a useful framework for identifying and interpreting functional dissociations among amygdala subregions in future studies.

A hemispheric analysis revealed that amygdala lateralization during negative affect was temporally restricted to the post-stimulus rating period, rather than the early stimulus-driven response. This pattern is consistent with prior evidence linking the left amygdala to sustained, cognitively mediated emotional processing ^29^, and may reflect processes such as sustained attention to internal affective states or memory encoding of emotional content that unfold after stimulus offset. Importantly, most prior neuroimaging studies of amygdala lateralization have focused on stimulus-evoked responses, which may explain why the literature presents an inconsistent picture ^39^.

Cognitive reappraisal did not substantially alter the early amygdala response in contrast to the robust temporal structure observed for passive stimulus viewing. Across both datasets, regulate and look-negative conditions produced highly similar evoked-response time courses during the initial picture-viewing window (approximately 3–6.5 TR), which is consistent with our previous work using canonical HRF model ^7^. Only weak and inconsistent differences emerged in the centromedial amygdala during the later rating period. Given the amygdala’s role in rapid and coarse recognition of emotional salience^40,41^, this pattern suggests that reappraisal does not intervene at the early period of appraisal to blunt the initial affective response. As a deliberate, effortful process dependent on lateral prefrontal control, reappraisal unfolds over several seconds ^42^, whereas the early amygdala response reflects rapid sensory-evaluative processing that is largely complete before top-down signals can take effect.

## Limitations

Several limitations of the present study warrant discussion. First, although FIR modeling allows flexible estimation of the temporal dynamics of the BOLD signal, the inferred neural dynamics remain constrained by the hemodynamic response. The amygdala is a small subcortical structure with vascular and physiological properties that may differ from neocortex ^43^, and even with FIR modeling, our temporal estimates cannot directly index neural firing.

Second, our subregion definitions were derived from probabilistic anatomical atlases, which approximate but do not perfectly correspond to cytoarchitectonic boundaries, and partial-volume effects at standard 3T resolution limit the precision with which small nuclei can be distinguished. Future work using ultra-high-field (7T) imaging may further refine the temporal characterization of amygdala subregions.

Third, both datasets used a TR of 2 seconds, which limits the precision with which fast affective dynamics can be resolved. Although the FIR analysis revealed a clear delayed centromedial response relative to laterobasal and superficial subregions, this temporal resolution does not support strong claims about causal ordering or directed interactions among amygdala nuclei.

Thus, the delayed CM response should be interpreted as a robust temporal dissociation, rather than evidence that earlier LB or SF activity causally drives later CM activation. Future work using faster acquisitions or complementary methods with higher temporal resolution will be needed to test causal models of information flow within the amygdala.

## Conclusions

Our findings show that the human amygdala is temporally heterogeneous during negative affective processing. Across two independent fMRI datasets, FIR modeling revealed dissociable response profiles across amygdala subregions: laterobasal and superficial subregions showed earlier, stimulus-locked responses, whereas the centromedial amygdala showed a delayed and more sustained response that extended into the post-stimulus evaluation period. This temporal dissociation was replicated across two anatomical atlases and was further supported by data-driven clustering of voxelwise time courses, indicating that temporal dynamics capture meaningful functional organization within the amygdala. By contrast, instructed cognitive reappraisal did not reliably alter these subregional BOLD time courses, suggesting that this form of regulation has limited influence on amygdala dynamics in the present task. Together, these results highlight the value of modeling temporal response structure and provide a more refined account of how amygdala subregions contribute to negative emotional processing.

## Methods

### Participants

Participants were drawn from two independent fMRI datasets collected as part of larger community-based health studies: the Adult Health and Behavior Project, Phase 2 (AHAB) and the Pittsburgh Imaging Project (PIP). Both studies recruited midlife adults from the Greater Pittsburgh area of Pennsylvania through mass mailings to residents of Allegheny County. Study 1 included 182 participants (95 males and 87 females; mean age = 43.45 ± 7.26 years) and Study 2 included 176 participants (87 males and 89 females; mean age = 40.58 ± 6.33 years). All participants provided written informed consent prior to participation. The University of Pittsburgh Human Research Protection Office approved both studies (Study 1 Protocol ID: 07040037; Study 2 Protocol ID: 07110287) as well as their aggregation into a common data registry (Protocol ID: 19030174).

Common exclusion criteria across both studies included self-reported history of diagnosed cardiovascular disease or clinical treatment for cardiovascular disease, psychotic disorder such as schizophrenia, chronic renal or hepatic disease, history of seizure or cerebrovascular disorder, ongoing or recent cancer treatment, chronic respiratory disease, and pregnancy.

Participants were also excluded for current use of medications to control insulin, glucose, glucocorticoids, arrhythmias, blood pressure, lipids, weight, or mood, and for standard MRI contraindications such as claustrophobia or metallic implants. Additional exclusion criteria specific to each study are detailed in ^7^

### Task and Paradigm

The task and paradigm are described in full in ^7^ and are summarized here. During fMRI scanning, participants completed an emotional picture viewing and cognitive reappraisal task. Each trial began with a 2-second instruction cue (’Look’ or ’Decrease’), followed by a 7-second image presentation period, a 4-second self-report rating period during which participants rated their negative emotion on a five-point scale (1 = neutral, 5 = extremely unpleasant), and a randomized rest period of 1 to 3 seconds. Participants viewed 30 unpleasant and 15 neutral images drawn from the International Affective Picture System (IAPS) ^44^. The task comprised 15 trials each of three conditions: ’Look neutral’, in which participants passively viewed neutral images; ’Look negative’, in which participants passively viewed unpleasant images; and ’Regulate negative’, in which participants were instructed to decrease their negative feelings toward unpleasant images by cognitively reappraising the circumstances and content depicted. Images were presented in pseudo-random order, with no more than two identical cues or four negative images presented consecutively. The task was administered using E-Prime software.

Normative arousal and valence values for images in Study 1 were comparable across ’Look negative’ (arousal: 6.16 ± 0.52, valence: 2.13 ± 0.37) and ’Regulate negative’ (arousal: 6.21 ± 0.67, valence: 2.01 ± 0.35) conditions, with neutral images substantially less aversive (arousal:2.65 ± 0.40, valence: 5.12 ± 0.57). Values in Study 2 were similarly matched: ’Look negative’ (arousal:6.14 ± 0.53, valence: 2.10 ± 0.37), ’Regulate negative’ (arousal:6.37 ± 0.52, valence: 2.02 ± 0.36), and ’Look neutral’ (arousal:3.44 ± 0.37, valence: 5.97 ± 0.35). Both studies used 1–9 scales to rate valence and arousal.

### MRI Acquisition

MRI data from both studies were collected on the same 3T Siemens Trio TIM whole-body scanner using a 12-channel phased-array head coil. Functional imaging parameters were as follows: field of view = 205 × 205 mm, matrix size = 64 × 64, TR = 2,000 ms, TE = 28 ms, and flip angle = 90 degrees. A high-resolution T1-weighted MPRAGE structural image was also acquired for each participant for purposes of co-registration and normalization.

### fMRI Preprocessing

Functional data were preprocessed using Statistical Parametric Mapping software (SPM12; http://www.fil.ion.ucl.ac.uk/spm). Slice timing correction was applied prior to realignment to account for acquisition timing differences across slices in this event-related design. Functional images were then realigned to the first image in the series using a six-parameter rigid body transformation. Realigned images were co-registered to each participant’s skull-stripped and bias-corrected MPRAGE T1 image, and subsequently normalized to Montreal Neurological Institute (MNI) space and interpolated to 2 × 2 × 2 mm voxels. Normalized images were spatially smoothed using a 6 mm Gaussian kernel.

Nuisance regressors included six head motion parameters (x, y, z, roll, pitch, and yaw), spike indicator regressors for potential motion outliers (volumes falling outside the 95% confidence region of the multivariate distribution of images in multidimensional space, defined by Mahalanobis distance), and extracted timeseries from cerebrospinal fluid masks using CANlab tools (https://github.com/canlab/CanlabCore). A high-pass filter of 1/180 Hz was applied to remove low-frequency temporal drift.

### Amygdala Atlas Definitions

Amygdala subregions were defined using two independent probabilistic atlases to ensure that findings were not dependent on any single parcellation scheme. The first atlas was the Amunts et al. cytoarchitectonic atlas ^24^, as implemented in the CANlab 2018 combined atlas (https://canlab.github.io), which defines three amygdala subregions: centromedial (CM), laterobasal (LB), and superficial (SF). The second atlas was the Nacewicz et al. atlas^25^, which parcellates the amygdala into four subregions: central and medial nuclei (CeMe), cortical and anterior amygdaloid area (CoAHA), basolateral and basomedial nuclei (BLBM), and the lateral nucleus (LA). Both atlases are available as part of the CANlab neuroimaging pattern masks repository (https://github.com/canlab/Neuroimaging_Pattern_Masks). Spatial smoothing was performed before ROI extraction; while this improves signal-to-noise ratio, it may also cause partial-volume mixing between adjacent subregions. Because such mixing homogenizes responses, our estimates are conservative.

### FIR Model Estimation

To characterize the temporal dynamics of the BOLD response without imposing assumptions about response shape, we applied a finite impulse response (FIR) modeling approach ^11,45^ at the first level. For each participant, -1 to 13TR relative to cue onsets for each condition (’Look neutral’, ’Look negative’, and ’Regulate negative’) were modeled using a set of delta functions spanning 15 successive TRs (30 seconds), yielding a freely estimated evoked-response time course of BOLD activity for each condition. This window was selected to encompass the full trial duration, including the cue period (1 TR), stimulus presentation (3.5 TRs), and rating period (2 TRs), as well as the subsequent hemodynamic return to baseline (∼6 - 7 TRs). Nuisance regressors were identical to those described above. Group-level FIR time courses for each subregion were estimated by averaging voxelwise FIR beta estimates within each atlas-defined region of interest across participants. Statistical significance of each time point relative to baseline was assessed using one-sample t-tests, with significance thresholded at p < 0.05 corrected for multiple comparisons across time points. We also conducted exploratory analyses using smooth FIR modeling ^45^. However, because the relatively short interval between trials in the current design increased the possibility that smoothed estimates would be influenced by temporally adjacent trials, we did not use the smooth FIR estimates in the primary analyses.

### Statistical analyses

All condition and region contrasts were conducted at the TR level rather than on pre-specified time windows. For each subregion, we tested (i) activation against baseline at each TR using one-sample t-tests; (ii) paired contrasts between conditions (look-negative vs. look-neutral; look-negative vs. regulate-negative) at each TR; and (iii) paired contrasts between the centromedial subregion (CM in Amunts atlas, CeMe in Nacewicz atlas) and each other subregion at each TR, testing where in time subregional divergence occurs. For each contrast, the p-values (one per TR) were corrected using the Benjamini-Hochberg FDR procedure at q < 0.05. Reported significant TRs are those surviving FDR correction within the corresponding contrast. For the descriptive onset claim (first TR at which look-negative BOLD rises reliably above baseline), we restrict the reporting window to TRs 3–13; earlier TRs fall within the hemodynamic pre-rise window and are not interpretable as onset estimates.

### Data-Driven Clustering of Amygdala Voxels

To test whether temporal dynamics alone could recover the anatomical organization of the amygdala, we performed a voxelwise clustering of FIR time courses using all three conditions jointly. Each voxel within the combined amygdala mask (the union of the Amunts and Nacewicz atlases, 958 voxels) contributed a 45-dimensional feature vector (15 TRs × 3 conditions) obtained by averaging the FIR estimates across the 358 participants. We clustered this feature matrix using k-means with squared Euclidean distance. To determine the optimal number of clusters, we applied a permutation-based model selection procedure ^46^. For each candidate k (ranging from 2 to 7), we computed the mean silhouette value, a measure of how well each voxel fits its assigned cluster relative to neighboring clusters, for both the observed data and 1,000 randomly permuted datasets in which each feature column was independently shuffled to destroy temporal structure while preserving marginal distributions. A pseudo-Z statistic was computed for each k as the standardized difference between the observed mean silhouette value and the null distribution mean: pseudo-Z = (observed silhouette − null mean) / null SD. The optimal k was selected as the value yielding the largest pseudo-Z, reflecting the greatest statistical separation between the true clustering solution and chance. This analysis indicated that a four-cluster solution was optimal under this criterion.

We assessed how well the data-driven clusters recovered each anatomical parcellation (Nacewicz, Amunts) using two indices, computed separately per atlas. Each atlas was first resampled into the cluster-map space, and all comparisons were restricted to voxels labeled in both maps . Both indices were tested against permutation nulls (10,000 iterations, fixed seed) built by shuffling cluster labels across the overlap voxels while holding anatomy fixed; because permutation only relabels voxels, this preserves cluster sizes and anatomical structure while removing any true correspondence.

Global agreement between the four-cluster and anatomical partitions was quantified with the Adjusted Rand Index ^47^ . Region-specific correspondence was quantified with the Dice coefficient, 2|C ∩ R| / (|C| + |R|), computed for every cluster–region pair; the full matrix of one-tailed p-values (4 × 4 for Nacewicz, 4 × 3 for Amunts) was corrected with Benjamini-Hochberg FDR at q < 0.05, and each cluster’s best match was the region with the highest Dice.

Because each permutation null is set by the sizes of the two voxel sets being compared, the significance of a Dice value depends on these sizes and not on its magnitude alone: chance overlap grows with set size, so a smaller Dice against a compact region can exceed its null while a numerically larger Dice against an extensive region may not.

## Data availability

The single-subject univariate beta images for both datasets are available through NeuroVault at https://neurovault.org/collections/16266/ (persistent identifier: https://identifiers.org/neurovault.collection:16266). The single-subject FIR estimates and source data underlying the figures and statistical analyses in the present study will be made publicly available upon publication.

## Code availability

MATLAB code and toolboxes used for the analyses are available through the CANlab GitHub repository at https://github.com/canlab. Custom code used for FIR estimation, amygdala subregion analyses, lateralization analyses, and data-driven clustering will be made publicly available at https://github.com/KeBo2018/KeBo2026_Amygdala_HRF upon publication.

## Acknowledgements

We thank S. Boyko and C. DuPont for assistance with data collection and scoring and Thomas E. Kraynak for contributions to participant recruitment and data preprocessing. This work was supported by the National Institutes of Health through grants R01MH076136 to T.D.W. and M.A.L and P01HL040962-25 to P.J.G. The funders had no role in study design, data collection and analysis, the decision to publish, or preparation of the manuscript.

## Supplementary Materials

**Supplementary Figure 1.**
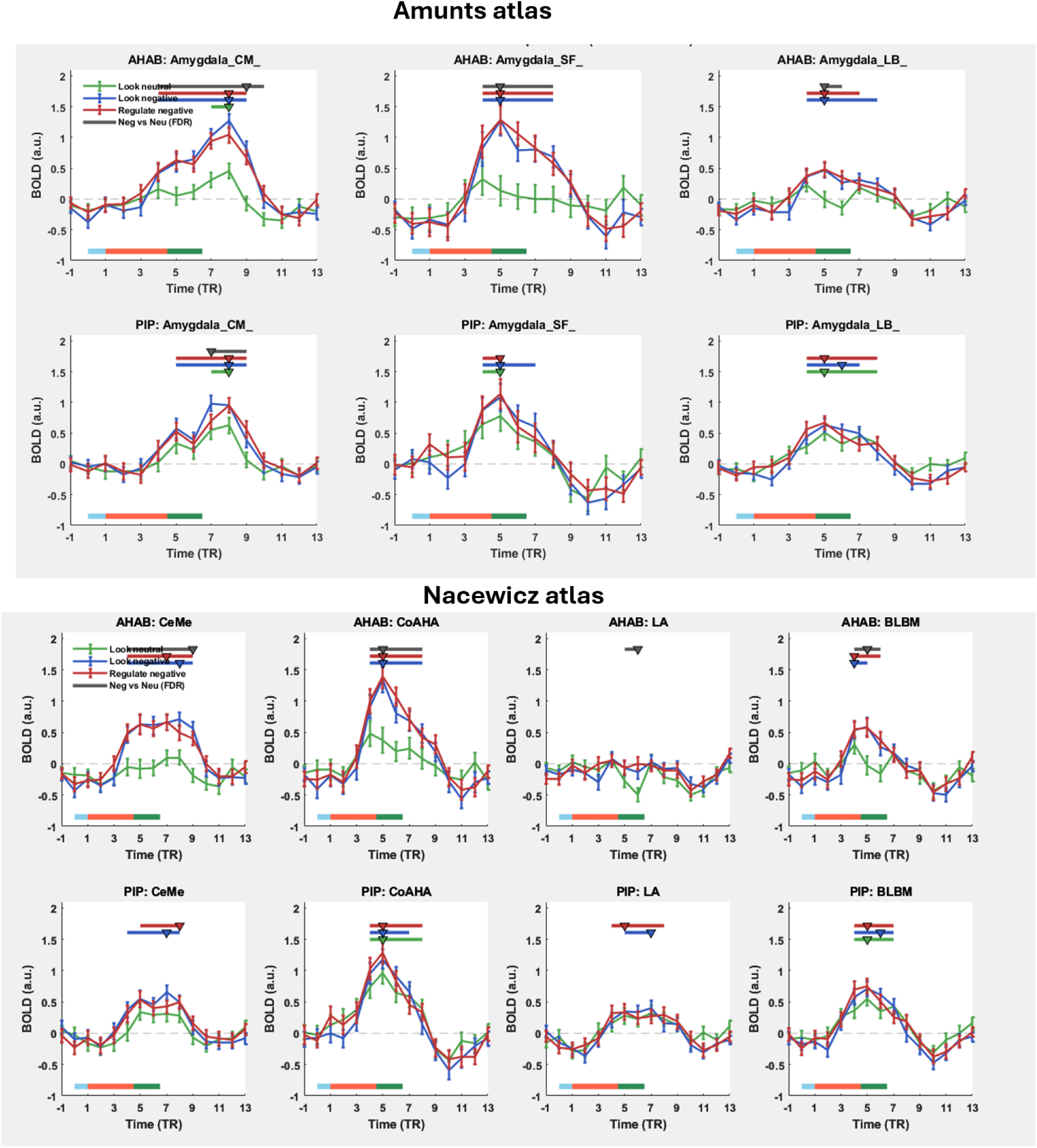
Temporal dynamics of amygdala subregions during emotional picture viewing and reappraisal, estimated by finite impulse response (FIR) modeling. BOLD responses were modeled with an FIR basis set spanning 30 s (15 TRs; TR = 2 s) for each of three task conditions. This analysis reproduces the pooled analysis shown in Figure 2 separately in the two independent cohorts. Region-level dynamics replicated across cohorts. The delayed response peaking after picture offset in CM amygdala reproduced in both cohorts. In the Look Negative condition, CM (Amunts) peaked at TR 8 in AHAB and TR 7 in PIP, and CeMe (Nacewicz) likewise at TR 8 and TR 7, whereas the superficial (SF), laterobasal (LB), cortical (CoAHA), and basolateral (BLBM) subregions peaked early (TR 5) in both samples; the lateral nucleus (LA) was the weakest and most variable region, as in the pooled analysis (peaking at TR 13 in AHAB but TR 7 in PIP, with near-zero AHAB amplitude, 0.09). Group-mean Look Negative evoked-response time courses were highly correlated across cohorts for every subregion except LA (Amunts: CM r = 0.95, SF r = 0.86, LB r = 0.93; Nacewicz: CeMe r = 0.91, CoAHA r = 0.91, BLBM r = 0.93; LA r = 0.57). The Look-Negative-versus-Look-Neutral differentiation of CM/CeMe reproduced robustly in AHAB (CM TRs 4–9; CeMe TRs 4–9) but only at isolated TRs in PIP (CM TRs 7 and 9; CeMe TR 4), while the corresponding differentiation of the superficial/laterobasal and CoAHA/BLBM subregions was clearer in AHAB than in PIP. As in the main analysis, reappraisal (Regulate vs Look Negative) produced no consistent, replicable modulation: the few TRs reaching the FDR threshold in AHAB (CM 3/8/13; CeMe 3/8/9) did not reappear in PIP (CM none; CeMe 7), so no region showed a reproducible reappraisal effect.

**Supplementary Figure 2.**
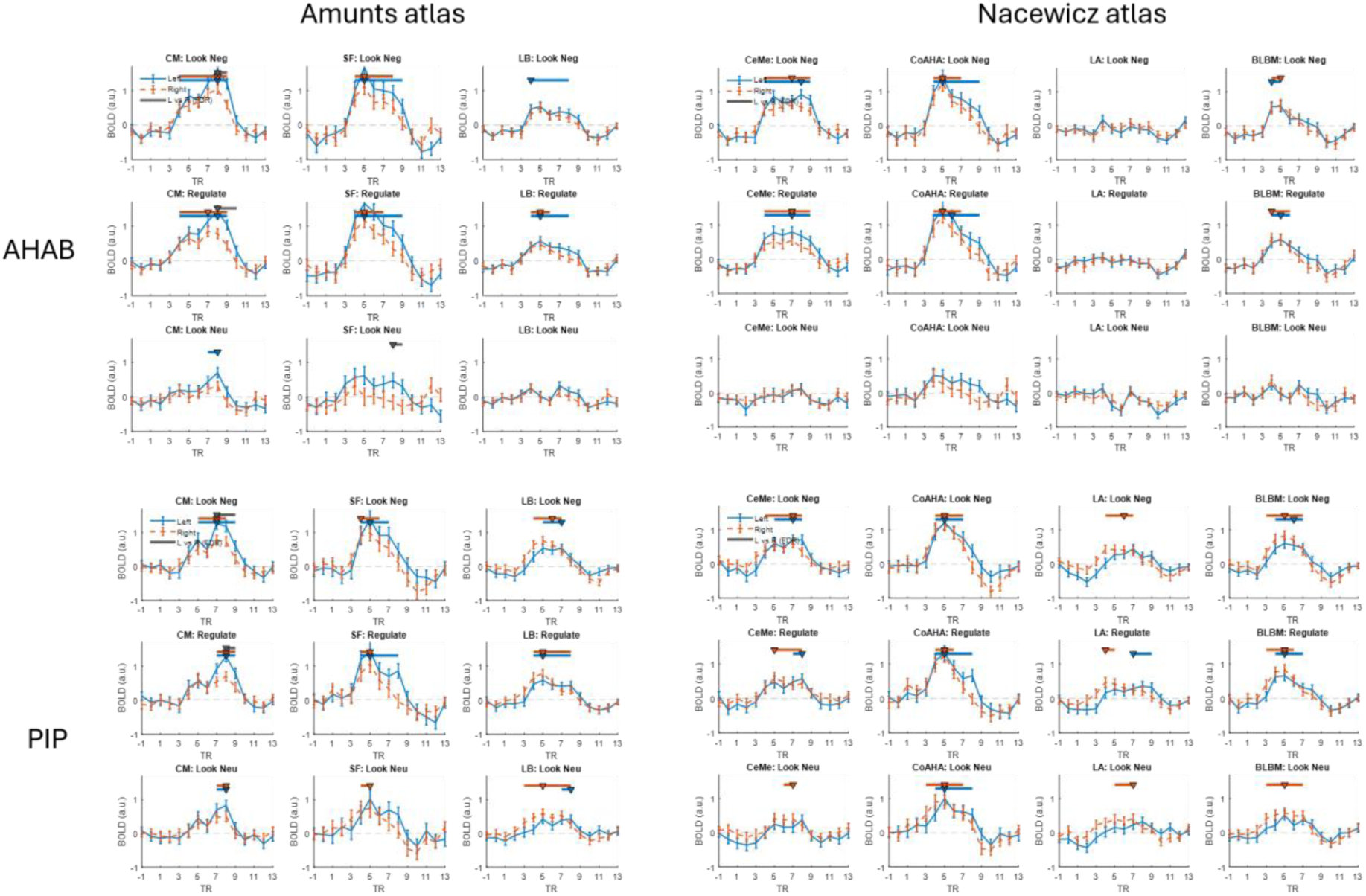
Hemispheric lateralization of FIR-estimated BOLD responses. Each panel shows group-mean FIR time courses (±SEM) for left (solid blue) and right (dashed red) hemisphere subregions across three conditions: look negative (top), regulate (middle), and look neutral (bottom). The x-axis represents time post-cue onset in TRs (TR = 2s); the y-axis represents BOLD amplitude in arbitrary units. Lateralization replicated selectively, in the centromedial subregion. The leftward asymmetry of the Amunts centromedial subregion during negative processing reproduced robustly in both cohorts (CM Look Negative LI: AHAB = +0.194, t = 4.63, p < .001; PIP = +0.162, t = 3.54, p < .001), both bracketing the pooled value of +0.187. The per-TR Left > Right effect likewise reproduced at late TRs in both samples for Look Negative (AHAB: TR 9; PIP: TRs 8–9) and Regulate (AHAB: TRs 9–10; PIP: TR 9), and was absent during Look Neutral in both, as in the pooled analysis. The smaller leftward asymmetry of the superficial subregion reached significance in AHAB (LI = +0.113, t = 2.35, p = .020) but not in PIP (LI = +0.042, p = .43), and the laterobasal subregion was non-lateralized in both cohorts (AHAB LI = +0.032, p = .46; PIP LI = −0.032, p = .52). For the Nacewicz atlas, no subregion showed a laterality index that was significant in both cohorts (CeMe: AHAB p = .076, PIP p = .49; CoAHA: p = .58 / .71; LA reached significance in AHAB only, p = .024 / .68), consistent with the main-text observation that no Nacewicz region was reliably lateralized. Thus the only lateralization effect to replicate across both independent samples was the leftward dominance of the centromedial amygdala — the same subregion carrying the late-sustained temporal profile.

**Supplementary Figure 3.**
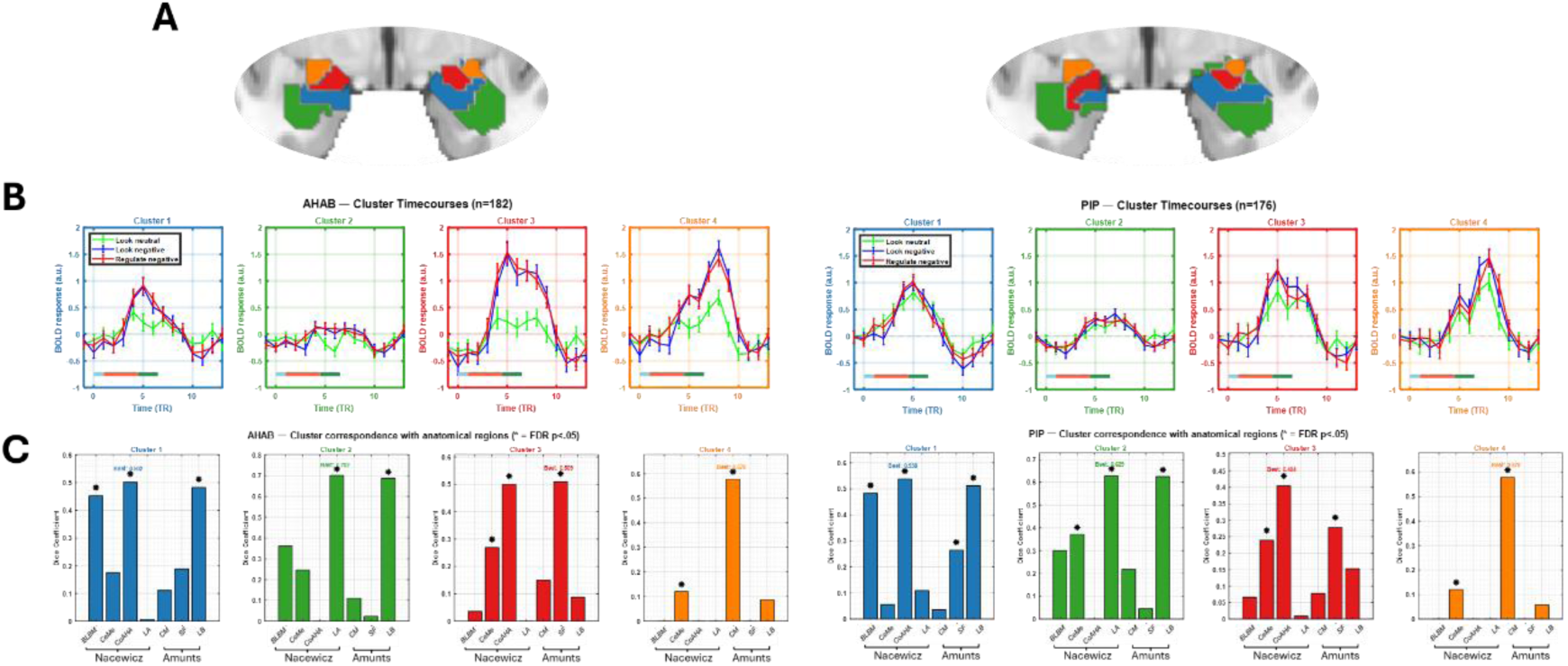
Replication between two datasets on clustering analysis: Clustering the 958 amygdala voxels on their 45-dimensional FIR feature vectors (k = 4) yielded comparable four-cluster partitions in each cohort. The data-driven partition corresponded to both anatomical atlases well above chance in each sample (AHAB: ARI = 0.258 [Nacewicz] and 0.210 [Amunts]; PIP: ARI = 0.214 and 0.132; all permutation p ≤ .0002), consistent with the pooled values reported in the main text (0.276 and 0.161). The four clusters were themselves reproducible across cohorts: the AHAB and PIP partitions agreed well above chance (ARI = 0.369, permutation p = .0001), and every AHAB cluster had a clear PIP counterpart (cross-cohort Dice = 0.62–0.79; mean best-match Dice = 0.71). Matched-cluster time courses were highly correlated across cohorts for the task conditions (Look Negative: mean r = 0.92, range 0.86–0.97; Regulate: mean r = 0.90), and lower for Look Neutral (mean r = 0.64), as expected given the minimal evoked signal in that condition. Critically, the late-sustained centromedial cluster replicated. In both cohorts, the cluster overlapping the Amunts CM mask did so at an identical Dice of 0.578 (matching the pooled value of 0.58); this same cluster showed the most sustained late Look-Negative-versus-Look-Neutral differentiation in each sample (AHAB: TRs 7–12; PIP: TRs 9–10); and the two cohorts’ centromedial clusters were matched to one another at the highest cross-cohort overlap of any pair (Dice = 0.791), with near-identical temporal profiles (Look Negative r = 0.971). Thus the late-sustained centromedial cluster identified in the pooled analysis emerged independently in each cohort.

